# Dynamic Attentional Control Through Mixed Prefrontal Cortex Resources

**DOI:** 10.1101/2024.10.03.614523

**Authors:** Fabio Di Bello, Francesco Ceccarelli, Adam Messinger, Aldo Genovesio

## Abstract

Spatial attention can be either involuntarily drawn to salient, unexpected events (exogenous attention) or voluntarily directed (endogenous attention). Evidence suggests a link of both forms of attention to the (oculo)motor system. We investigated the role of the prefrontal cortex (PFC) in this relationship using a task that spatially dissociated attention from motor planning in two monkeys. PFC units flexibly encoded either the attended location or motor target, with some showing overlapping coding. Endogenous attention promotes an early engagement of these shared resources, producing two key effects: (1) it prevents interference between motor and attention systems, and (2) it regulates the extent of the exogenous shifts for attentional capture, with stronger shifts occurring when endogenous attention is weak. Overall, we show that there are separate attention and motor systems, each of which can flexibly draw on some “overlapping” resources based on environmental and internal demands.

## Introduction

In our daily lives, we often observe a tight coupling between spatial attention and the oculomotor system, with rapid eye movements frequently accompanying shifts of attention. However, attention can also shift covertly, without an accompanying eye movement. Furthermore, attention can be directed either voluntarily based on internal goals (endogenous attention) or involuntarily drawn to salient or unexpected sensory events in the environment (exogenous attention). Endogenous attention is also known as top-down or goal-directed attention, while exogenous attention is also known as bottom-up or stimulus-driven attention. The interest in the relationship between covert attention and motor planning grew in the ‘80s, following the proposal that spatial attention directly results from motor planning (Klein R.M, 1980; Rizzolatti et al., 1987). According to this “premotor theory of attention”, a unified system (the motor system) is responsible for both motor planning and attention. This theory predicts that, if the motor system is compromised, both endogenous and exogenous attention should be impaired. However, studies with neuropsychological patients with damaged oculomotor control have shown deficits specific to exogenous attention, while endogenous attention remained largely unaltered (Gabay et al., 2010; Rafal et al., 1988; Sereno et al., 2006; Smith et al., 2004). Using an eye abduction protocol that compromised saccade execution, Smith et al. (2012) found behavioral deficits in exogenous attention while endogenous attention remained unaffected, indicating that only reflexive attention relies on oculomotor preparation. These different relationships with the motor system may indicate that endogenous and exogenous attention operates through different mechanisms (Chica et al., 2013; Landry et al., 2021; Pinto et al., 2013; Xia et al., 2024). However, others have found a link between them, with endogenous attention modulating exogenous attention (Keefe et al., 2021; Meyer et al., 2018). For instance, endogenous attention can prevent attentional capture of an unexpected distractor when oriented elsewhere (Di Bello et al., 2022; Theeuwes, 1991). Also, endogenous attention enhances the effect of exogenous attention when they co-localize (Dubey et al., 2023; Müller & Rabbitt, 1989; Theeuwes, 2014). The independence or not between these two attentional mechanisms has important neurofunctional implications, as it indicates whether separate or partially overlapping brain regions are involved (Chica et al., 2013). To date, the interplay between these two types of attention and their relationship with the motor system remains an open question in ongoing research (Lowet et al., 2018; Michalczyk et al., 2020; Sapountzis et al., 2022).

To address this, we investigated the contributions of the prefrontal cortex (PFC), a region central to both attentional control and oculomotor processes (Bowling et al., 2020; Fernández et al., 2023; Martinez-Trujillo, 2022), to the interplay between covert attention and oculomotor dynamics in monkeys performing a task that spatially dissociated these variables. The task included cue and uncued trials. Cued trials involved endogenously attending to one of four possible locations (the attention target) to detect a subtle brightening (the Go-signal) that signaled when to make a saccade to a distinct oculomotor target location. In uncued trials (20% of trials) only the motor target was specified, making the location of the Go-signal unpredictable and detection of the peripheral brightening a function of exogenous attention.

Consistent with our prior research (Messinger et al., 2021), we have identified neural units solely dedicated to motor or attention processes, contradicting the assumptions of the premotor theory of attention. However, we also found that some units exhibited flexibility, serving both functions. We refer to the latter as “overlapping” resources. In uncued trials, when endogenous attention was not directed to a specific target, the overlapping resources were more involved in motor planning, enhancing motor target representation but causing interference when attention later shifted to the Go-signal. In contrast, during cued trials, the endogenous deployment of attention coordinated these resources more effectively, allowing motor and attention processing to function independently, thereby reducing interference between the two systems. Using time-resolved cross-modal neural decoding, we found that saccade planning and attention share a similar spatial representation, both before and after the attention target presentation. Notably, classifiers trained to decode the motor plan were also effective at decoding exogenous attention shifts, the extent of which was influenced by the degree of endogenous attention deployment.

Our findings support the notion that the endogenous and exogenous systems are independent yet can interact under certain conditions. We found evidence that a partial PFC overlap between the motor and attentional systems provides the neural basis for this interaction. This allows for a dynamic balance between internal needs and external demands, similar to flexible buffer systems (Fiebelkorn & Kastner, 2020; Carrasco, 2011; Chica et al., 2013).

## Results

### Behavior

Monkeys R and G performed the behavioral task shown in Fig. 1 correctly on 80% and 81% of trials, respectively. Error trials were mostly caused by a lack of behavioral response, suggesting that the Go-signal at the Attended Location went undetected. Such saccade omissions occurred in 17% of trials for Monkey R and 16% for Monkey G. Saccades to the wrong motor target were much less frequent, occurring in only 3% of trials for Monkey R and 2% for Monkey G. The incorrect saccades were equally distributed amongst the non-targets for Monkey R, whereas they were disproportionately directed toward the Go-signal location for Monkey G (32% and 52% of incorrect saccades, respectively). Task accuracy on cued trials, where the location of the Go-signal was signaled beforehand, was significantly greater than on uncued trials (Monkey R: 81.5 ± 1.1% versus 75.2 ± 1.6%, 36 sessions, Kruskal-Wallis nonparametric test, P < 0.001; Monkey G: 83.8 ± 0.5% versus 73.6 ± 1.0%, 42 sessions, Kruskal-Wallis nonparametric test, P < 0.001). Reaction times (RTs) were also faster on cued trials compared to uncued trials (Monkey R: 411 ± 3.4 ms versus 419 ± 3.2 ms, 36 sessions, Kruskal-Wallis nonparametric test, P < 0.012; Monkey G: 313 ± 1.6 ms versus 323 ± 2.3 ms, 42 sessions, Kruskal-Wallis nonparametric test, p < 0.001). These results suggest that both monkeys used the cue’s center color to covertly attend to the region of space where the Go-signal would be presented.

**Figure 1.**
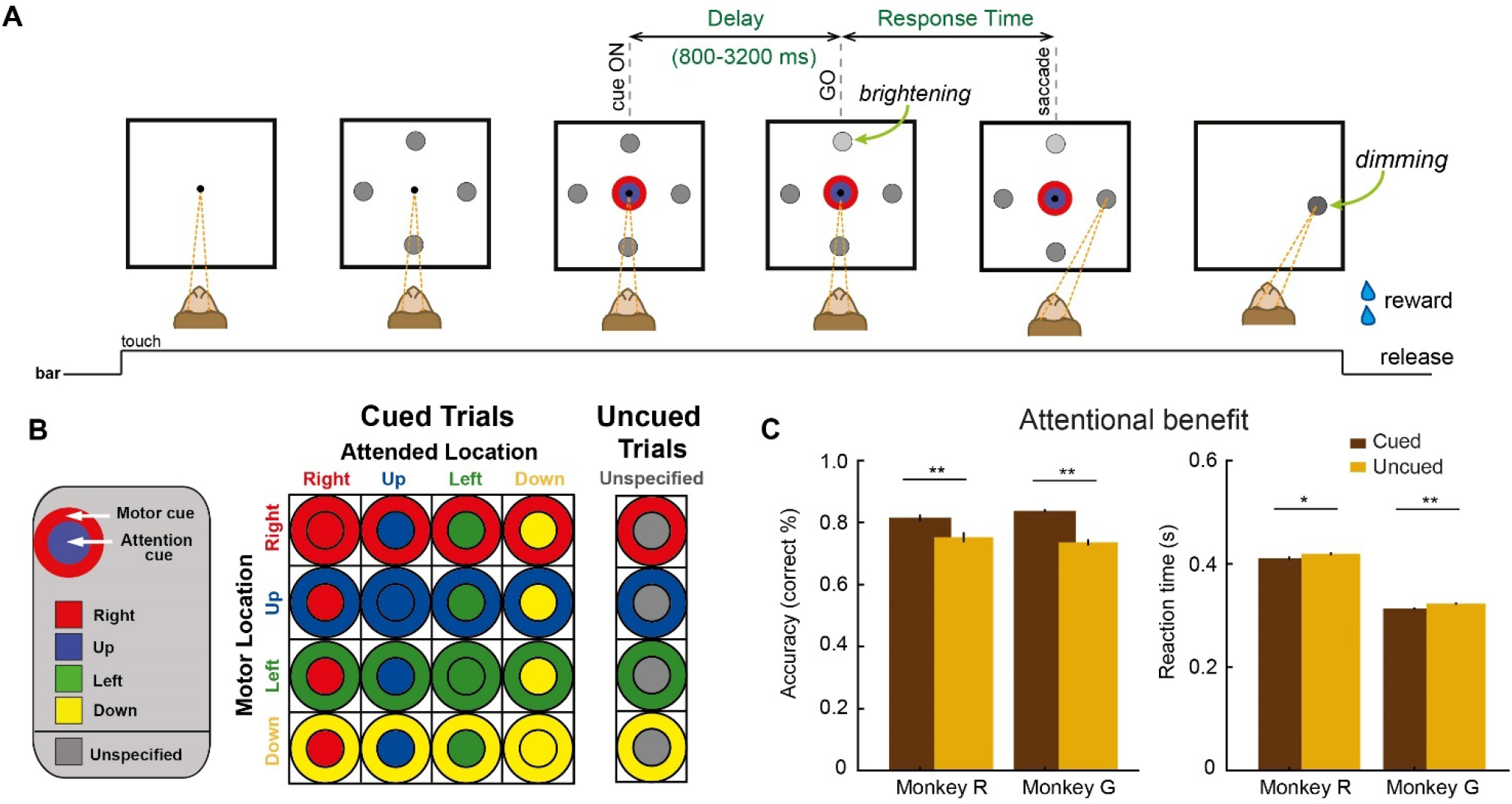
Behavioral task and cueing effect. (**A**) The monkey (shown from above) pressed a bar to make a central fixation spot appear. After fixation (convergent dashed lines) four peripheral gray landmarks were presented, followed by a two-part colored cue stimulus centered on the fixation point. After a variable cue delay period, a subtle brightening (the Go-signal) of one of the peripheral spots indicated to the monkey when to direct its gaze to a particular target spot. The cue’s inner color indicated the Go-signal location: the attention cue. The cue’s outer color specified the target of the upcoming saccade: the motor cue. In the example highlighted in (A and B) the cue’s blue inner color signifies that the Go-signal would be located up (the attended target), and the cue’s red outer color specified a saccade to the rightward gray spot (the motor target). (**B**) The stimuli that served as cues for the different combinations of Motor and Attended Locations are represented in a 4 × 4 matrix. On uncued trials (20% of correct trials), the inner circle was gray and the Go-signal location was selected randomly so the monkey did not know where to attend. The motor target was still specified by the color of the annulus. (**C**) Behavioral results: Percent of trials in which each monkey succeeded to detect the Go-signal and complete a saccade (left panel) and the average Reaction Time (right panel), for cued (brown bars) and uncued (light brown bars) correct trials. Error bars: SEM across sessions. *p < 0.05, **p < 0.001.

### Prefrontal neurons are modulated by attention and by oculomotor targets

We tested whether the prefrontal cortex encoded the attention target and motor target specified by the central cue stimulus. A two-way ANOVA was used to assess neuronal firing rates over a sliding window for 209 individual units, each recorded for three or more trials across the cued and uncued spatial conditions (as illustrated in the 4×4 and 4×1 matrices in Fig. 1B). Units significantly tuned for the motor target emerged ∼250ms after cue onset and increased in number until ∼550ms (Fig. 2A, upper panels). Significant tuning for the Attended Location or for both the Motor Location and the Attended Location (hereafter referred to as ‘Both’ units) was less common, with these proportions increasing ∼400ms after cue onset (Fig. 2A, upper panels). As expected, on uncued trials (Fig. 2A, lower panels), significant attention tuning was largely absent during the delay period because the target of covert attention was not specified. Thus, PFC activity reflected oculomotor plans as well as whether and where covert attention was directed by the cue.

**Figure 2.**
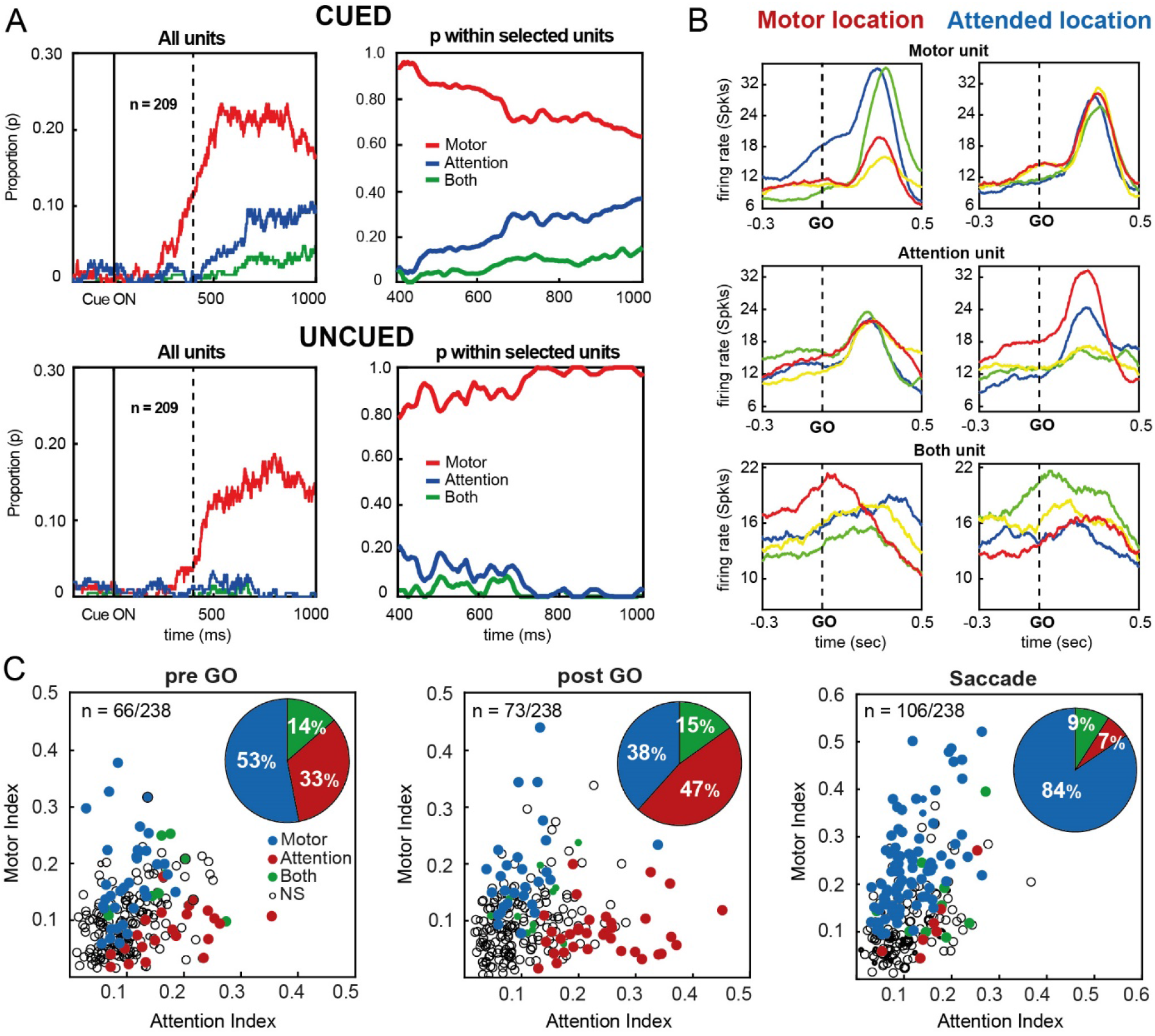
Temporal evolution of attention and motor selectivity in PFC. (**A**) Motor (blue lines), Attention (red lines) and Both (green lines) tuned units following cue onset (two-way ANOVA over a sliding window, p < 0.01) in cued (top panels) and uncued (bottom panels) trials as a proportion of all units (left panels) and of selective units (right panels). The time axes in the right panels begin at 400 ms, corresponding to the dashed lines in the left panels. (**B**) Neural tuning following the Go-signal (t=0) on cued trials for the four Motor Locations (left column) and the four Attended Locations (right column) of three example multi-units (rows) modulated by the motor target (Motor unit), the locus of attention (Attention unit), and both locations (Both unit), respectively. (**C**) Scatter plots showing the motor versus attention modulation indices for all recorded units on cued trials (N = 344), calculated based on firing rates in the pre-Go (−200 to 0 ms before the Go-signal), post-Go (100 to 300 ms after the Go-signal), and post-Saccade (600 to 800 ms after the Go-signal) epochs. Units were classified during each epoch based on significant tuning for the Attended Location (Attention), Motor Location (Motor), both the Attended and Motor Locations (Both), or neither variable (not significant, NS). Pie charts show the proportions of significant units of each type in each epoch. Larger symbols in the Pre-Go scatter plot correspond to the three units shown in (B).

It has been shown that PFC shows a diverse range of responses for various cognitive variables in different task epochs (Amengual et al., 2022; Friedman & Robbins, 2022; Rigotti et al., 2013). To investigate this in our task, we analyzed how PFC units were modulated by motor and attention variables in cued trials, using the 238 units recorded for at least three trials in each of the 16 possible spatial conditions (see the 4×4 matrix in Fig. 1B). Fig. 2B illustrates examples of Motor, Attention, and Both units before and after presentation of the GO signal. Of the units exhibiting selectivity before the Go signal (the ‘pre-Go’ epoch), 53% were Motor units, 33% were Attention units, and 14% were Both units (see Fig. 2C and Table S1). The breakdown of selectivity shifted in the period after the brightening of the attention target (the ‘post-Go’ epoch), with the proportions of Attention units increasing to 47%, Motor units decreasing to 38%, and the Both units remaining stable at 15%. In the period after the motor response (the ‘post-Saccade’ epoch), there was a jump in the proportion of Motor units (84%) and a drop in Attention and Both units (7 and 9%, respectively). Overall, the number of significantly selective units gradually increased, with 66 units in the pre-Go epoch, 73 in the post-Go epoch, and 106 in the post-saccade epoch.

Table S1 illustrates the evolution of unit selectivity over time, including units classified as untuned. Some initially untuned units developed Attention (9.3%), Motor (8.6%), or Both (4.6%) selectivity following the Go-signal, while after the saccade, many untuned units developed Motor selectivity (Motor = 27.9%; Attention = 6.4%; Both = 8.1%). Units that were motor selective (i.e., Motor and Both Units) in the pre-Go epoch mostly retained their motor selectivity in the post-Go (50%) and post-saccade (77%) epochs. Most units that were initially attention selective (i.e., Attention and Both Units) retained their attention selectivity into the post-Go epoch (52%) but not the post-saccade (13%) epoch. Other units encoded a different single variable between epochs, without ever simultaneously encoding both variables. For example, three Motor units from the pre-Go epoch switched to encoding only the Attended Location in the post-Go epoch and nine units initially classified as Attention units became only motor target selective in the post-Saccade epoch (Table S1).

### Decoding of decoupled motor and attentional information

We employed a decoding technique to compare the PFC’s parallel coding of attention and motor variables during both cued and uncued trials (see Methods), focusing on critical epochs (i.e., around Cue onset, Go-signal presentation, and Reward delivery). To better assess the independent encoding of each variable in cued trials, we focused only on trials for which the Attended and Motor Locations differed (i.e., “different” trials, off-diagonal conditions in Fig. 1B).

#### Motor target information dynamic

Following the Cue onset (Fig. 3, upper-left panel), we observed above chance decoding of motor information starting at 250 ms and 200 ms for cued and uncued trials, respectively (Kruskal-Wallis nonparametric test, p < 0.001, Cue onset aligned). This significant motor decoding accuracy persisted throughout the delay period, with uncued trials showing higher motor accuracies than cued trials from 400 ms onward. This enhanced classification accuracy on uncued trials was still present at the end of the delay period (−150 ms to -100 ms relative to the Go-signal, Fig.3, upper-middle panel).

**Figure 3.**
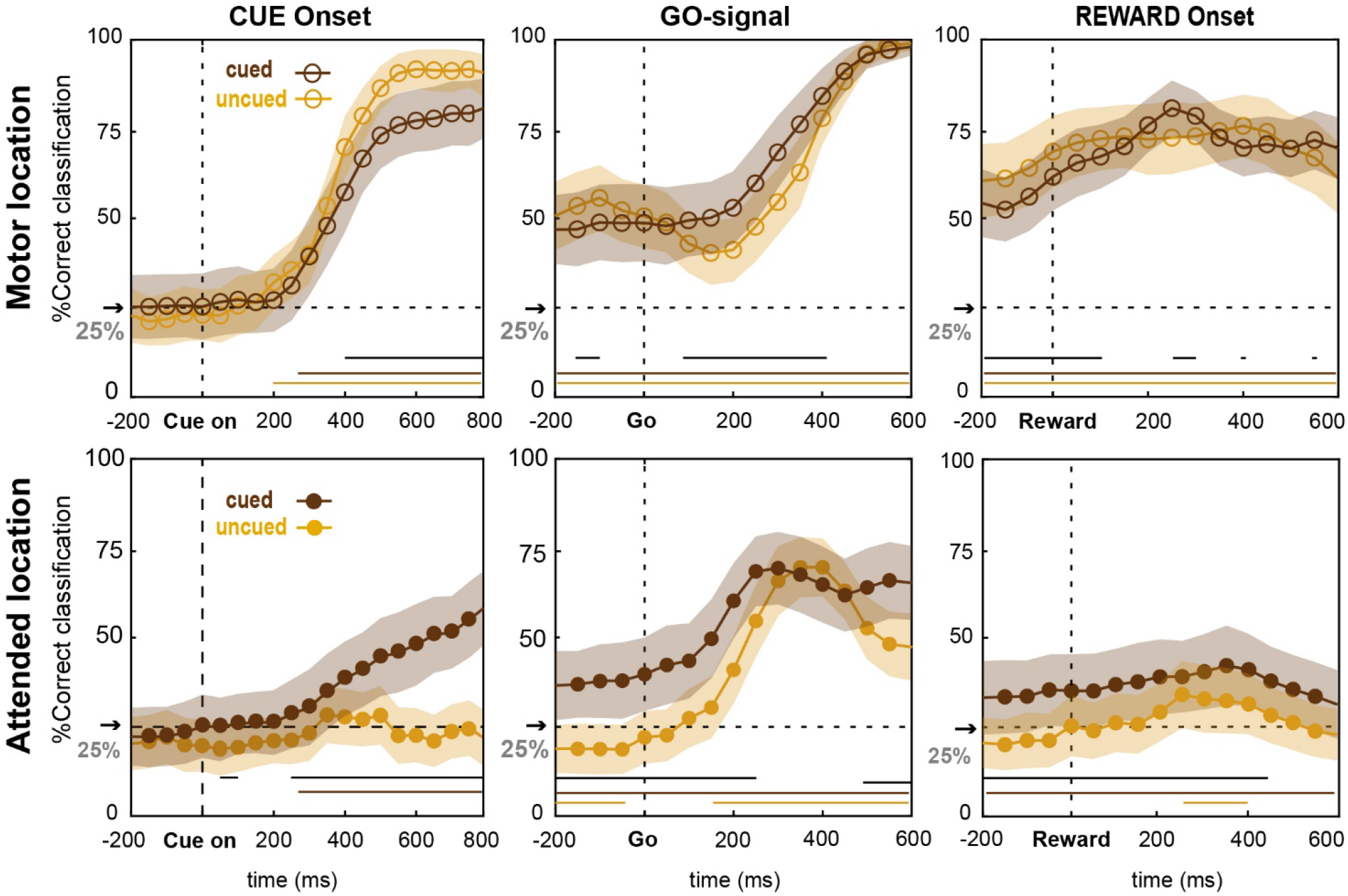
Population decoding of movement and attention during three task periods. Time course of population decoding accuracy for the Motor Location (top row) and Attended Location (bottom row) on cued (dark brown) and uncued (light brown) trials, aligned to three critical task events (Cue onset, Go-signal and Reward delivery). Lines and shading show the mean classification accuracy ± s.e. in each sliding window. Black horizontal lines indicate a significant difference between cued and uncued decoding accuracies (p < 0.001). Colored horizontal lines show significant (p < 0.001) deviation from chance (dashed horizontal line) for cued (dark brown) and uncued (light brown) trials.

Following the presentation of the Go-signal, the accuracy for decoding the Motor Location on uncued trials dipped to 40.7 ± 9.0% at ∼150 ms, which was significantly (Kruskal-Wallis nonparametric test, p < 0.001) less than at the time of the Go-signal (51.8 ± 9.2%). In contrast, on cued trial the decoding accuracy did not differ between these same time points (49.8 ± 10.6% at ∼150ms vs. 49.6 ± 10.4% at Go-signal time, Kruskal-Wallis nonparametric test, p = 0.96). Motor decoding accuracy on uncued trials was significantly worse than on cued trials from 100-400 ms after the Go-signal, even as decoding accuracy on both trial types improved to near perfect classification by 500 ms. Classification of the motor target remained well above chance on cued and uncued trials through reward delivery and beyond (Fig.3, upper-right panel). Classification accuracy was significantly higher on uncued trials than cued trials until 100 ms after reward delivery, after which there were no sustained accuracy differences.

In summary, PFC represented the motor target from saccade planning until long after saccade completion and reward delivery. Decoding accuracy was significantly different on cued and uncued trials despite the fact that the motor target instruction was present on both types of trials. Surprisingly, specifying where to covertly attend on cued trials resulted in a less accurate representation of the motor plan during the delay and less interference from the Go-signal in decoding the motor target, relative to uncued trials.

#### Attended target information dynamic

In cued trials, we observed significantly above chance decoding of the attention location starting ∼250 ms after Cue onset, with accuracy climbing throughout the delay period (Fig. 3, bottom-left panel). As expected, decoding of the Attended Location on uncued trials never exceeded chance level during the delay period and even dropped below chance late in the delay period. Later in the trial, the classification accuracy for the attended location in the uncued trials dropped below chance before the attended target appeared (from - 200 ms to -50 ms relative to the Go-signal).

Following the Go-signal presentation (Fig. 3, bottom-middle panel), we observed a sharp increase in decoding accuracy for the Attended Location in both trial types, peaking at approximately 300 ms in the cued trials and 400 ms in the uncued trials. We interpret these sudden increases in accuracy for the Attended Location as indicative of successful attentional capture of the attended stimulus (see discussion). Interestingly, these peaks reached a similar level of accuracy for classifying the Attended Location (cued = 69.4 ± 10.1% vs uncued = 69.6 ± 8.4%, Kruskal-Wallis nonparametric test, p = 0.71), suggesting a comparable amount of visual information was needed for detection on both trial types.

Decoding of the Attended Location remained accurate following the saccade (Fig. S1, bottom panel) on cued trials but decreased on uncued trials, with the two conditions significantly diverging starting 200 ms after the saccade. On cued trials, decoding of the Attended Location remained significantly above chance past the time of reward delivery (Fig. 3, bottom-right panel).

### Interaction between motor and attentional neural codes

The previous section described parallel motor and attention information coding on cued trials. Here, we explore the stability and the generalization of the coding for motor and attention across the epochs surrounding the brightening of the attended target (i.e., the Go-signal). For this, we adopted a cross-temporal decoding analysis (see Methods), in which a linear classifier was trained with spatial variables (motor target or attended target variables) using data in one period and tested with data from another period. Both “same” and “different” trials were taken into account (Fig. 1B).

#### Cross-temporal decoding within modalities – stability

Figure 4 shows the results of cross-temporal decoding when a single variable, either the Motor Location (A) or the Attended Location (B), was used for both the training and test phases of classification. The diagonal of the heat maps (Fig. 4AB, top panels) shows classifier performance when trained and tested on the same time bin. Accuracy along the diagonal is plotted in the ‘Diagonal performance’ panel below each heat map and was significantly above chance for all time bins (Kruskal-Wallis nonparametric test, p < 0.001). The time course of the accuracy along the diagonal is similar to that of cued trials in the preceding section (Fig. 3, middle panels), but with greater accuracy due to the inclusion of “same” trials. Fig. S2 shows that the accuracies shown in Fig. 4 were significantly greater with “same” trials included than without them (Kruskal-Wallis nonparametric test, p < 0.001 for all time bins from -200 to 400, step 50 ms, Go-signal aligned).

**Figure 4.**
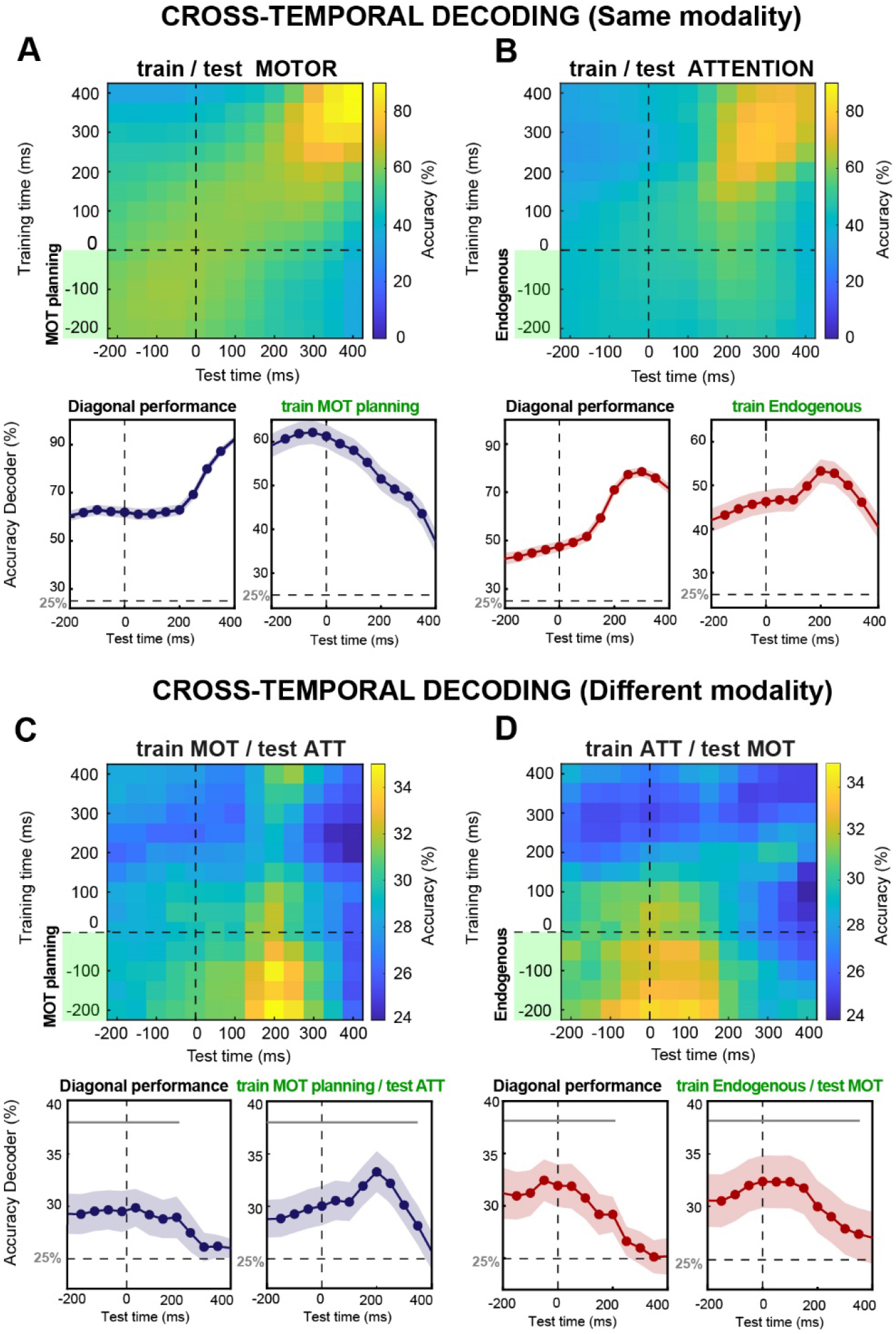
Dynamics and interplay of motor and attention coding around the Go-signal. (**A-B**) Cross-temporal classification for the Motor (**A**) and Attended (**B**) Locations. Heat map shows mean decoding accuracies over n = 250 resamples. The training time of the classifier is shown on the ordinate and the test time is displayed on the abscissa, both relative to the Go-signal brightening. The green shaded bars on the ordinate correspond to the training time interval used for the decoding shown in the bottom-right panels. For both motor and attention classification, average accuracies were calculated along the diagonal (bottom-left panels) and when training was restricted to the delay epoch only (bottom-right panels). All classifications in the bottom panels were significantly (p < 0.001) above chance (dashed horizontal line). (**C-D**) Cross-modal classification accuracies when the classifier was trained on the Motor Location and the Attended Location was decoded (**C**) and when trained on the Attended Location and the Motor Location was decoded (**D**). Bottom row, same conventions as in (A-B) but for cross-modal classification. Horizontal gray lines in bottom panels indicate time periods when classification was significantly (p < 0.001) above chance.

We observed differences in the stability of the motor and attention spatial representations across task epochs. Specifically, we examined how well a classifier trained on delay period activity (green shading on the y-axis of Fig. 4AB heat maps) decoded locations from activity following the Go-signal. We found that the motor planning code, which was predictive during the delay, lost its capacity to decode the motor target once the Go-signal was issued (Fig. 4A, bottom-right panel). In contrast, the delay period representation of endogenous attention continued to code the Attended Location after the Go-signal and even exhibited a significant improvement in classification that peaked at ∼200 ms (Fig. 4B, bottom-right panel; Accuracy at Go = 46.4 ± 1.6% vs. Accuracy at peak = 53.3 ± 1.7%, Kruskal-Wallis nonparametric test, p < 0.001).

#### Cross-temporal decoding across different modalities – generalizability

We next investigated how well each of the spatial task variables was decoded by a classifier trained on the other task variable (Fig. 4CD). This allowed us to assess the extent to which the neural representation of the motor target generalized to the encoding of attention and vice versa. We evaluated both these cross-modal tests along the diagonal (i.e., with matching train and test time windows) and found significant encoding of the tested variables throughout the delay epoch and continuing for 200 ms after the appearance of the Go-signal (Fig. 4CD, bottom panels).

The procedure for the cross-temporal analysis between modalities mirrored that for within modality analysis. A classifier trained on the motor target variable (i.e., motor planning activity) generalized to decoding the Attended Location during the delay period and the period around the Go-signal (Fig. 4C, bottom-right panel). This decoding, while less accurate than within-modality decoding (Kruskal-Wallis nonparametric test, p < 0.001 for all time bins from -200 to 400, Go-signal aligned), was significantly above chance. The accuracy of decoding of the Attended Location increased abruptly around 200 ms after the Go-signal (Accuracy at Go = 30.3 ± 1.5% vs. Accuracy at peak = 33.7 ± 1.6%, Kruskal-Wallis nonparametric test, p < 0.001). This suggests that the representation of space for oculomotor planning generalized to the localization of attention, and especially to the capture of exogenous attention by the Go-signal brightening. Decoding remained above chance until ∼350 ms.

For the reverse cross-modal analysis, we trained the classifier on delay period activity associated with the attended target (i.e., endogenous attention activity) and assessed classification of the motor target variable (Fig. 4D, bottom-right panel). Significant prediction of the motor target was found within the delay and for approximately 200 ms following the Go-signal, after which decoding accuracies gradually declined to chance level. This suggests that the spatial encoding of endogenous attention bares some similarity to that for motor planning, but differs from that for motor execution.

### Neuronal interplay between endogenous and exogenous attention

We have shown that behavioral performance (Fig. 1C) and neural classification of the Attended Location (Fig. 3) were better on cued trials than on uncued trials. By extension, we predicted that behavioral performance would be better when endogenous attention was precisely allocated to the Attended Location. To assess this, we estimated the location of the attentional spotlight (AS) and used its displacement (dAS) from the Attended Location to quantify the fidelity of covert attention to the cued attention target. Consistent with our expectation, we found (Fig. S3) that the attentional spotlight was significantly closer to the Attended Location (i.e., smaller dAS) on correct trials than on saccade omission trials, when the Go-signal was not detected. Our finding accords with previous studies that assessed attention through signal detection (Amengual et al., 2022; De Sousa et al., 2021; Di Bello et al., 2022).

To get insight into the neural dynamics underlying endogenous attention, we grouped trials into short (< 4.0°), intermediate (between 4.5° and 6.0°), and long (> 6.5°) dAS groups based on the proximity of endogenous attention to the Attended Location at the time of the Go-signal. We restricted analysis to trials where, at the expected time of Go-signal detection (∼250 ms after the Go-signal, see Methods), decoded attention was near the Go-signal location (dAS < 3°). On average, 42.2% of correct trials met this criterion. Motor decoding along the diagonal was largely similar across the three dAS groups (Fig. 5B, left panel). In contrast, attention decoding along the diagonal had substantial differences across the three dAS groups (Fig. 5C, left panel). Accuracy at the Go-signal was highest for short-dAS trials (90.8 ± 1.4%) and progressively lower as endogenous attention deviated further from the attention target (intermediate-dAS = 75.2 ± 2.3%; long-dAS = 44.1 ± 2.6% trials). This inverse relationship between dAS and decoder performance was present throughout the delay period and until 200 ms after the Go-signal, after which all three categories had the same high levels of classification accuracy.

**Figure 5.**
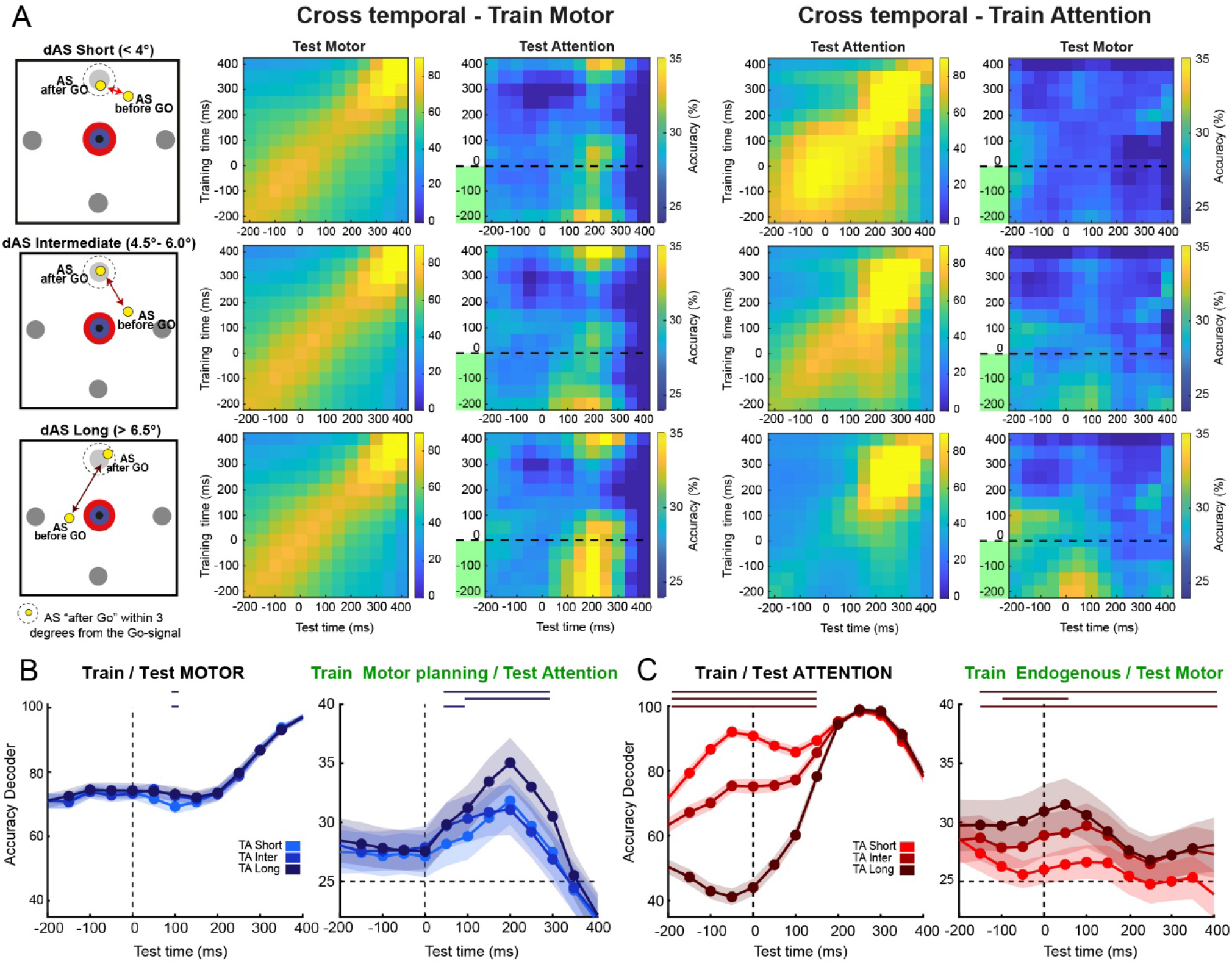
Interplay of neural codes as a function of attention localization. (**A**) Colormaps show decoding accuracy of cross-temporal classifiers trained on the motor code (2^nd^ and 3^rd^ columns) and on the attention code (4^th^ and 5^th^ columns) for within (2^nd^ and 4^th^ columns) and across (3^rd^ and 5^th^ columns) modality decoding. As depicted in the 1^st^ column, rows show analyses for trial with short, intermediate, and long displacements of attention (dAS) from the cued Attended Location, respectively. The dAS is the estimated misalignment of the attentional spotlight (AS) before and after the Go-signal occurs. (**B-C**) Left panels show the average within-modality decoding accuracy of cross-temporal classifiers trained and tested along the diagonal using the motor (attention) code derived from short-, intermediate-, and long-dAS trials. Right panels show the average cross-modal decoding accuracy for classifiers trained during the delay period (as indicated by the green shading on the y-axes of the 3rd and 5th columns in (A)) using short-, intermediate-, and long-dAS trials. Horizontal lines, from top to bottom, indicate time periods of significantly different decoding accuracy for long- vs short-dAS trials, long- vs intermediate-dAS trials, and intermediate- vs short-dAS trials, respectively (Kruskal-Wallis nonparametric test, p < 0.05).

Unlike “within modality” decoding, cross-modal decoding was more accurate when the AS was *further* from the attention target. Decoding of the Attended Location by classifiers trained on motor planning activity (Fig. 5B, right panel), exhibited an increase in accuracy following the Go-signal that peaked at ∼200 ms. The accuracy increase was progressively larger the further endogenous attention was from the cued attention target at the time of the Go-signal (long-dAS accuracies > short-dAS accuracies from 50 ms to 300 ms; long-dAS accuracies > intermediate-dAS accuracies from 150 ms to 300 ms; intermediate-dAS accuracies > short-dAS accuracies from 50 ms to 150 ms; Kruskal-Wallis nonparametric test, p < 0.05). This improvement in attention decoding suggests that the attention representation was most like that of motor planning when large shifts in exogenous attention were required to capture attention.

The other cross-modal classifier, trained on the endogenous attention signal and tested on decoding of the motor target, also performed better on trials where the endogenous AS was estimated to be further from the Attended Location (Fig. 5C, right panel. Long-dAS accuracies > short-dAS accuracies from -150 ms to 400 ms; long-dAS accuracies > intermediate-dAS accuracies from -100 ms to 50 ms; intermediate-dAS accuracies > short-dAS accuracies from -150 ms to 400 ms; Kruskal-Wallis nonparametric test, p < 0.05). For short-dAS trials, cross-modal decoding was not significantly different than chance, indicating that when attention was precisely allocated, there was little overlap in the neural representations of attention and motor planning. For long- and intermediate-dAS trials, however, accuracies were better than chance during the delay period and for the first 150 ms following the Go-signal (Kruskal-Wallis nonparametric test, p < 0.05) before gradually declining to chance level before the saccadic response. Thus, the less attention focused on the Go-signal location during the delay epoch, the greater the degree of cross-talk between the neural resources involved in saccade planning (but not execution).

Overall, our results indicate a partial overlap between the attentional and motor systems, enabling the use of exogenous attention when endogenous attention has not been accurately allocated to the attended position.

## Discussion

Recent evidence indicates that voluntary (endogenous) attention is largely independent of the motor system (Messinger et al., 2021) while involuntary reflexive (exogenous) attention is strongly connected to it system (Gabay et al., 2010; Lowet et al., 2018); implying the existence of two distinct types of attention (Chica et al., 2013). However, other studies indicate a connection between the two, with endogenous attention modulating exogenous attention (Keefe et al., 2021; Meyer et al., 2018). If these are indeed two independent systems that sometimes interact, we would expect them to be implemented in partially overlapping brain regions. However, evidence supporting such an overlap is still lacking.

In this study, we aimed to determine the PFC contribution to these two attentional modes during visual selection and their reciprocal relationship with the motor system. To address this question, we analyzed neural activity in the prefrontal cortex of two monkeys performing a behavioral task that spatially dissociated motor targets from the target of endogenous attention.

Briefly, we found that monkeys could use a centrally fixated cue to endogenously direct attention to a peripheral location that was distinct from their planned saccade target (Fig. 1). Endogenous attention accelerated detection of the Go-signal both in terms of behavioral reaction times (Fig. 1C) and neural decoding of this peripheral brightening event (Fig. 3, bottom). Neuronally, most units encoded either the locus of attention or the motor target, with relatively few units encoding both variable (Fig. 2). Tuning changed over the course of the trial, with attention tuning being most common from 400 ms following cue presentation until Go-signal detection. Directing voluntary attention away from the saccade target on cued trial impaired decoding of the motor plan but also made it resilient against a drop in decoding that occurred on uncued trial, when both the timing and location of the Go-signal were unpredictable (Fig. 3, top and S2). Cross-temporal and cross-modal decoding revealed further interplay between the attention and motor representations. A classifier trained to decode the motor target during the delay period (saccade planning stage) was not effective at decoding this variable around the time of saccade execution. However, it was effective at decoding where the Go-signal occurred, especially on those trials where we estimated that endogenous attention had strayed from its target (Fig. 4 and 5). In contrast, a classifier trained to decode the Attended Location based on delay period activity (i.e., an endogenous attention signal) showed improved decoding before the Go-signal for both attention (within modality) and the motor target (across modalities), again especially on trials where attention was off-target (Fig. 4 and 5).

### Overlapping motor and attention information in PFC

In our original study on these data, we showed that the delay period activity of single units in the prefrontal cortex (PFC) encoded either the oculomotor plan or covert spatial attention, but not both (Messinger et al., 2021). Here we have extended this finding to multiple task periods using the larger pool of PFC multi-units recorded on cued and uncued trials. This consistent finding that motor planning and spatial attention signals are encoded by distinct neural populations refutes the notion of obligatory coupling between these processes. This dissociation of functions at the neural level lies in contrast to the “premotor theory of attention”, which postulates that spatial attention is functionally equivalent to planning eye movements (Rizzolatti et al., 1987; Sheliga et al., 1994). Instead, we find that the PFC treats each of these cognitive operations distinctly.

Though most units were exclusively tuned for the motor or attention location, we found significant and rapid shifts in the relative proportion of such units across different phases of the task. Motor units outnumbered attention units early, during the delay period. Attention units became more numerous later, following the brightening of the attended target (the Go-signal). Finally, the predominance of motor units re-emerged around the time of the saccade. The proportion of ‘Both’ units remained almost unchanged throughout these events. Interestingly, units sometimes switched between motor and attention selectivity across task periods, in some cases without first simultaneously encoding both variables (i.e., without becoming ‘Both’ units in the interim). Thus, in any given epoch, largely separate neural units encoded the motor and attention variables of the task but the makeup of these two pools flexibly updated to reflect changing task demands throughout the trial, with some units serving both cognitive operations in different contexts. We refer to these units as ‘overlapping’ resources for their ability to be tuned for both variables across the trial. Previous studies have demonstrated that the PFC can encode and recruit different variables interchangeably depending on task demands (Genovesio et al., 2016; Kim et al., 2011; Marcos & Genovesio, 2016; Rougier et al., 2005). Here, we found that such flexibility extends to the interplay between motor and attentional systems.

### PFC dynamically recruits overlapping resources based on task demands

Our results indicate that PFC simultaneously represents both planned saccades and the targets of attention using motor selective cells and attention selective cells, respectively (Messinger et al., 2021). The analysis in Fig. 3 shows the decoding of these two variables on cued and uncued trials from a population of PFC units. Both locations were accurately and independently decoded, despite the fact that only trials with a distinct motor and attention target were used in this analysis.

On uncued trials, the location of the Go-signal was unpredictable and its detection had to rely on exogenous attention. We found that motor decoding was better in these trials than in cued trials when endogenous attention was engaged. Thus, when there were concurrent demands on the motor and attention systems, less motor plan information was encoded in the PFC. We infer from this result that when endogenous attention is not being taxed, the motor system recruits all available resources (both solely motor and the ‘overlapping’ resources) to encode the motor parameters, not because these extra resources are strictly required for solving the task, but simply because they are available (Lavie, 1995). This hypothesis gains support from two other observations. Firstly, we observed that the Go-signal resulted in a drop in motor decoding accuracy on uncued only. We attribute this drop in motor accuracy to interference produced by the exogenous attentional demands of the Go-signal brightening. Consistent with our hypothesis, this interference suggests the motor system was utilizing ‘overlapping’ resources on uncued trials that were suddenly reclaimed by the attention system to process the unpredictable Go-signal. Secondly, we found that just before the Go-signal, attention decoding on uncued trials was not at chance (as expected) but below chance. On uncued trials, the ‘overlapping’ resources that normally participate in attention decoding instead represent the motor target. Since for this analysis of ‘different’ trials, the motor target was never the same as the attended target, the stronger representation of the motor target comes at the expense of even chance classification of the attended location. The suggested neural dynamics imply a high capacity for neural flexibility within the PFC, where units can encode either information based on encountered needs. As mentioned earlier, some PFC units had such neural flexibility.

In contrast to uncued trials, in the cued condition, the sudden increase of attentional information in response to the attended target did not interfere with motor coding, supporting the idea that ‘overlapping’ resources have already been allocated and/or distributed between the systems before the presentation of the attended target (Corbetta, 1998; Kowler et al., 1995). Unlike the uncued condition, we observed that units encoding the Attended Location maintained their high level of coding even after the attended target appearance. Next, we will discuss the hypothesis of a mechanism for routing neural resources into endogenous attention. Here, we want to introduce the possibility that an early allocation of overlapping resources might leave traces of its ‘route’ (i.e., the neural connections of the endogenous neural pathway) even after target detection, thereby maintaining high predictions in the subsequent epochs. However, greater stability of spatial representations may also depend on the involvement of mnemonic processes during endogenous attention. For instance, it is known that spatial attention can rely on functional markers for location-specific representations held in working memory, thereby providing an additional contribution to sustained neural representation without necessitating a shift in resource encoding (Awh & Jonides, 2001; Panichello & Buschman, 2021; Smyth & Scholey, 1994). Further in-depth investigations are required to disambiguate this issue.

### Interaction between endogenous and exogenous attention mechanisms

As discussed above, our results suggest a significant role for overlapping neural resources in exogenous attention. This form of reflexive attention revealed itself through a sharp increase in decoding accuracy following the appearance of a salient or unexpected stimulus (Hunt et al., 2019). This sudden increase in the processing of attentional information results from the rapid allocation of attentional focus toward the stimulus, a phenomenon commonly known as ‘attentional capture (Luck et al., 2021; Theeuwes, 2014).

Although stimulus-driven attentional capture has been mainly studied in the context of the distracting impact of irrelevant targets on visual search tasks (Folk et al., 1992; Theeuwes, 1992), this phenomenon also occurs for selecting relevant information, and this happens even during the active employment of spatial attention (Di Bello et al., 2022). In line with this, we observed a sudden increase in attentional information in response to target stimuli in cued trials, revealing an attentional capture mechanism during endogenous attention (Fig. 3). Our cross-temporal analysis (Fig. 4B) indicates that the endogenous code contributes to this capture, as shown by the rapid increase in predictiveness following the Go-signal. Although this may challenge the exclusive association of attentional capture with exogenous attention, one might argue that the endogenous component of attentional capture can also make use of overlapping resources recruited during the delay period (rather than being used to code the motor information or not engaged at all). Thus, the reflexive component of attentional capture during endogenous attention may primarily depend on the involvement of these resources. This hypothesis is supported by the analysis shown in Fig. 4C, where we found that a classifier trained on the motor planning signal exhibited a sudden improvement in decoding the attention location, even in the presence of endogenous attention. This implies that 1) overlapping resources encoding motor variables during the delay period are employed in exogenous attention after the Go-signal and that 2) exogenous attention co-occurs with endogenous attention. This is consistent with our previous observation that exogenous attention relies on the rapid and extensive recruitment of overlapping resources, as well as with the well-established link between exogenous attention and the motor system (Gabay et al., 2010; Lowet et al., 2018). According to our classification, these overlapping resources are primarily visuomotor units (Dubey et al., 2023; Moore et al., 2003), which facilitate faster redirection of attention toward a salient stimulus compared to purely visual units. We speculate that the latter may be more involved in the slower process of endogenous attention by enhancing the receptive field’s sensitivity toward the attended target location (Desimone & Duncan, 1995; Reynolds & Heeger, 2009).

Recent evidence suggests that voluntary attention deployment fluctuates over time. Spatially and temporally resolved decoding of the attentional spotlight’s locus has revealed its dynamic nature, which is not strictly confined to the cued location (Di Bello et al., 2022; Fiebelkorn & Kastner, 2020; Gaillard et al., 2020). In this study, we observed that the perception of a low-saliency target-stimulus depended on the position of the attentional spotlight just before its presentation. By investigating the involvement of overlapping resources in relation to the endogenous orienting we found that 1) the mutual contribution of overlapping resources in attentional capture depended on spatial attention: the more distant the attentional spotlight, the more pronounced their involvement and vice versa. Moreover, 2) a focused orienting of attention significantly reduced the endogenous code’s ability to predict the motor target. We interpret that, in this condition, overlapping resources were heavily recruited to support spatial attention, leaving limited capacity for motor location coding. This is consistent with our observation that in conditions of less focused orienting, the endogenous code’s ability to predict the motor target gradually increased as the distance of the attentional spotlight from the attended target increased. These findings support the idea that overlapping resources are in a push-pull relationship with attentional orienting: the less attentional orienting, the greater the overlapping resources participate in the capture process.

Overall, our data support the notion that endogenous and exogenous attention are independent mechanisms that can interact under certain circumstances. As Chica et al. (2013) suggested, “Two independent systems that sometimes interact are expected to be implemented in partially overlapping brain regions.” We have demonstrated that in the PFC, a partial overlap between the motor and attentional systems may form the neural basis through which exogenous and endogenous attentional modes interact.

### Motor and Attention as interconnected but separate systems

The relationship between spatial attention and the motor system has been a longstanding debate in neuroscience. Early studies suggest a significant overlap between these systems. For example, electrical microstimulation of frontal regions has been shown to affect both oculomotor planning and endogenous attention (Beckers et al., 1992; Grosbras & Paus, 2002; Moore & Armstrong, 2003; Moore & Fallah, 2001; Muri et al., 1996; Thickbroom et al., 1996). While compelling, these findings are complicated by the possibility that microstimulation may engage neighboring but distinct neural networks simultaneously (Tehovnik, 1996), making it difficult to determine their independent functional contributions. Similarly, human neuroimaging studies suggest that preparing eye movements to a specific location activates the same frontal and parietal cortical networks involved in covert attention (Beauchamp et al., 2001; Corbetta, 1998; De Haan et al., 2008; Nobre et al., 2000; Perry & Zeki, 2000). However, the limited spatial and temporal resolution of fMRI makes it difficult to distinguish between signals from nearby but distinct neural circuits, such as those in the PFC. On the other hand, other studies challenge the idea of such an extensive overlap, suggesting that attention and eye movements rely on largely separate systems. This view is supported by evidence showing distinct neural populations in the PFC responsible for visual selection and saccade programming (Juan et al., 2004; Messinger et al., 2021; Murthy et al., 2001; Sato & Schall, 2003; Thompson et al., 1997, 2005).

An emerging alternative view reconciles these perspectives by proposing that motor and attention systems are distinct but sometimes interconnected, drawing on limited shared resources when needed (Fiebelkorn & Kastner, 2020; Hunt et al., 2019; Sapountzis et al., 2022). These systems often function independently, especially during endogenous attention, but become more coupled during exogenous attention. Our study aligns with this perspective. We found that the motor and attention systems exhibited a high degree of functional independence. However, we also observed a partial overlap between these systems. This overlap was indicated by (1) a small proportion of neural units in each task period that were simultaneously tuned to both spatial variables and (2) cross-modal decoding analyses, which demonstrated that the delay-period representation of the motor plan and the allocation of attention carried information relevant to predicting the other variable. To our knowledge, this is the first direct evidence of a partial overlap between these two systems in the PFC based on the refined spatial and temporal resolution provided by electrophysiological signals. These ‘overlapping’ resources can flexibly support either system depending on the available information, predominantly processing motor variables in uncued trials and both motor and attentional variables in cued trials. We provide evidence of the key role of these overlapping resources in attentional capture, with their involvement contingent upon endogenous attention deployment.

The presence of overlapping resources between the motor planning and attention systems does not directly support the predictions of the “premotor theory of attention,” which posits that endogenous attention uniquely emerges as a direct consequence of subthreshold motor planning. Our results align more with the idea that while distinct, the motor and attentional systems can leverage partially overlapping resources to flexibly balance environmental circumstances and internal needs.

## Material and Methods

All procedures were performed in accordance with the Guide for the Care and Use of Laboratory Animals and were approved in advance by the Animal Care and Use Committee of the National Institute of Mental Health.

### Behavioral task

We trained two male rhesus monkeys (Macaca mulatta), monkey R (9.0 kg) and monkey G (7.5 kg), to perform the task illustrated in Fig. 1A. Each monkey sat in a primate chair facing a video screen 57 cm away with its head fixed by a surgically implanted head post. The experimental design dissociated the locus of covert spatial attention from the target of a future saccadic eye movement. To accomplish this objective, we used a dual conditional visuospatial task. One four-way conditional association provided the spatial goal for a saccade; another four-way conditional association guided the allocation of covert attention. The monkey began each trial by touching a metal bar, which led to the appearance of a small white circle (0.15° radius), the fixation point, at the center of the video screen (Fig. 1A). If the monkey continued to fixate this spot [within a square window of 3.3° for a variable period (200 to 1000 ms)], four circular gray spots appeared (0.35° radius): left, right, up, and down from the fixation point at an eccentricity of 4.0°. A bicolored cue stimulus appeared at the fixation point after an additional fixation period of 200 to 1000 ms. The color of the cue’s outer component, a 0.25°-wide annulus, informed the monkey about which of the four gray spots to fixate after a forthcoming Go-signal. For convenience, we call this spot the Motor Location and its instructing stimulus the motor cue. The color of the cue’s inner component, a 0.6° radius circle, indicated which of the four gray spots would subsequently brighten as the Go-signal. We call this spot the Attended Location, and its central instructing stimulus is the attention cue. By design, the monkey should direct covert attention to the Attended Location to facilitate detection of the brightening event, which triggered a saccade to the Motor Location. The degree of brightening was selected randomly among four calibrated levels such that each monkey failed to respond to the smallest degree of brightening on ∼30% of trials and responded reliably to the largest degree of brightening. We refer to the variable and randomly selected interval (800, 1600, 2400, or 3200 ms) between the appearance of the cue stimulus and the Go-signal as the cue delay period. During this interval, the monkey should direct covert attention to the spot at the Attended Location while simultaneously planning a saccade to the spot at the (usually distinct) Motor Location. Following the Go-signal, the monkey had to complete a saccade to the Motor Location within 800 ms (for monkey R) or 600 ms (for monkey G). If it did so, then all other stimuli disappeared from the screen, and the monkey had to fixate the Motor Location (within a square window of 4.0°) for an additional 800 to 1600 ms until it dimmed. The monkey could then release the bar to receive a juice reward. If the monkey broke fixation before the Go-signal, failed to make a saccade within the requisite time after the Go-signal, shifted fixation anywhere other than to the Motor Location, failed to fixate the spot at the Motor Location for the required interval, or released the bar before this spot dimmed, the trial ended without reward delivery, and the normal interval between trials (600 to 1800 ms) was extended by 1600 ms. The same conditional mapping between the four cue stimulus colors and the four peripheral spots was used for the motor and attention cues. As illustrated in Fig. 1B, red was associated with the gray spot to the right of the fixation point, blue with up, green with left, and yellow with down. This arbitrary mapping was overlearned by trial and error before the neuronal recordings. In 20% of the trials, the colored attention cue was omitted and replaced with a gray cue of the same dimensions. On these uncued trials—i.e., uncued with respect to covert attention—the location of the Go-signal was selected pseudorandomly from amongst the four peripheral spots.

### Surgery and neuronal recording

A metal head post was surgically implanted under anesthesia before training began. After each monkey had learned the task, we surgically implanted a 27 × 36 mm recording chamber over the left frontal cortex, with its long side oriented in an anterior-posterior (AP) plane and its short side in a medial-lateral (ML) plane. A craniotomy was centered at AP +29.5, ML +12 in monkey R and AP +22, ML +15 in monkey G. Recordings were made in broad frontal regions, (see Fig. 5 in Messinger et al., 2021 for a map of the recording sites) PF included parts of the dorsolateral and ventrolateral PF on either side of the principal sulcus (areas 46d/v, 8, and 45). Recordings in this area were made using a multielectrode microdrive, with independently moveable single-contact electrodes arranged in a circle. The initial penetrations in both monkeys were made with a 7-electrode System Eckhorn drive (Thomas Recording GmbH, Giessen, Germany). Later penetrations were made with Alpha Omega’s 8-electrode MultiDrive (Alpharetta, GA). Neighboring electrodes were 1.0 mm apart, and the largest electrode separation. MUA signal was extracted using an auto threshold method. The threshold was set at negative 3.5 times the standard deviation.

### Modulation index calculation

For each unit, and for both motor and attention spatial variables, the modulation index was defined as MI = (maxFR – minFR) / (maxFR + minFR) where maxFR and minFR correspond to the higher and the lower mean firing rate across trial emerging within the 4 [UP, LEFT, DOWN, RIGHT] spatial conditions. Firing rates were computed on the calculated based on firing rates in the pre-Go (−200 to 0 ms before the Go-signal), post-Go (100 to 300 ms after the Go-signal), and post-Saccade (600 to 800 ms after the Go-signal) epochs.

### Motor and attention encoding in cued and uncued trials

To explore motor and attention encoding dynamics in both cued and uncued trials, we applied a decoding analysis on trials where the motor and attention target locations were dissociated (i.e., ‘different’ trials). To evaluate the classification accuracy with distinct randomly chosen trials, we executed 250 resample runs. In each run, we included all units recorded in at least 8 trials for each of the four possible motor or attention target locations. This created a matrix of 214 units by 32 trials (8 trials per motor or attention target location) for both cued and uncued conditions.

For decoding, we employed a leave-one-trial-out methodology. In this approach, one trial is designated the test set, and (nearly) all other trials form the training set. To avoid imbalance in the training set, three trials were removed in addition to the test trial such that all locations were equally represented in the training set. During the training phase, for each time step, a regularized optimal linear estimator (RegOLE) (Di Bello et al., 2021; Gaillard et al., 2020; Amengual et al., 2022) associated the neural responses, consisting of a vector containing the neural signals collected at each recording contact, from a structure containing the signal of 28 correct trials (i.e., 7 trials for each target location) to the (x,y) coordinates of the associated target position for each trial. To avoid over-fitting, we used a Tikhonov regularization, which gives us the following minimization equation: norm (W∗(R+b)−C)+λ∗norm(W). The scaling factor λ was chosen to allow for a good compromise between learning and generalization (De Sousa et al., 2021). Specifically, the decoder was constructed using two independent regularized linear regressions, one classifying the x-axis (two possible classes: −1 or +1) and one classifying the y-axis (two possible classes: −1 or +1). During testing, the classifier’s output was estimated from a vector of the average neuronal activity for each of the 214 recorded channels for the time interval of interest on a testing trial, new to the classifier. If the classifier’s output fell within the same quadrant as the Go-signal, the classification was considered correct. Classification accuracy was the average proportion of correctly classified trials within each resample run.

We used a nonparametric random permutation approach to statistically evaluate the significance of decoding accuracies to estimate the 99.9% confidence interval limit. We randomly reassigned each trial’s behavioral classification and recalculated the decoding accuracy. This process was iterated 250 times, generating a null distribution of accuracy values representing chance performance. The accuracy of the real non-permuted data was considered significantly above chance if it fell within the 0.1% upper tail of its spatially defined random permutation distribution.

### Cross-Temporal Decoding Procedure in cued trials

To explore the stability of attentional and motor codes around the time of the Go-signal on cued trials, we employed a cross-temporal decoding analysis. In this method, a classifier was trained with one spatial variable (the motor target or attended target variable) using data from one period and then tested with data from another period. To ensure unbiased classifications, for each channel, we randomly selected 10 correct trials for each of the 16 types of cued trials (comprising both ‘same’ and ‘different’ trials) formed by the 4 × 4 combinations of motor and attended targets (Fig. 1B). This yielded a matrix of 216 units by 160 trials. The classifier was then trained on 80% of the trials and tested on the remaining 20%, constituting a previously unseen set of trials. We performed 250 resample runs to evaluate the classification accuracy with distinct sets of randomly chosen trials. Due to the symmetry of the matrix and the simultaneous task instruction, we used the same data to train both the motor and attention classifier on each run. By utilizing the same classifier described earlier (the RegOLE), we associated the neural responses from a dataset comprising signals from 128 correct trials (8 trials for each combination of target locations) with the (x,y) coordinates of the corresponding target positions for each trial for each time-bin. We then tested this association using a dataset for the time interval of interest consisting of 32 testing trials (i.e., 2 trials for each combination of target locations), which was new to the classifier. Classification accuracy was the average proportion of correctly classified trials within each resample run. Thus, by training and testing the classifier using all possible combinations of 50ms, we obtained a classification accuracy matrix where the values along the diagonal were calculated by training and testing on equivalent time bins. In contrast, different time bins were used to calculate the off-diagonal values.

A cross-modal decoding approach was also applied to investigate the generalizability of the motor and attentional codes. We used the same methodology described for cross-temporal decoding. However, in this analysis, the type of variable associated with training and the one being tested belonged to different modalities (i.e., motor and attention target locations were alternately used for training and testing).

### Target to decoded Attentional Spotlight distance computations

The (x,y) readout of the attention decoder was not categorically assigned to one of the four possible Attended Locations as is commonly done with classifiers, but was taken as a continuous measure of the position of the attentional spotlight (AS) (Di Bello et al., 2021; Ameungual et al., 2022). Here (Fig. S3) we show that this estimated readout of the AS position accounts for variations in behavioral responses.

For each trial and each attention location, we measured the Cartesian distance (dAS) between the decoded AS at Go-signal onset and at the expected perceptual seizure (i.e., at 250 ms – as inferred by Figure 3 and Figure 4) to determine if deviations in the locus of attention were reflected in neuronal MUA responses.

We performed 250 resample runs to evaluate dAS distances with distinct randomly chosen trials to collect a large dataset of dAS-associated trials. For each run, we generated a matrix of 216 units by 160 trials by randomly selecting 10 correct trials for each of the 16 spatial cue types. We categorized the fidelity of attention localization on extracted trials as high, intermediate or low based on whether the deviation of the attentional spotlight was short (dAS < 4.0°), Intermediate (4.5° < dAS < 6.0°), or Long (dAS > 6.5°). We used these trial categories in cross-temporal decoding analyses, as previously described, except that the input matrix consisted of neuronal activity on trials belonging to the same dAS category.

## Acknowledgments

We thank Dr. Steve Wise for his numerous contributions and Dr. Andrew Mitz for engineering. We thank also K. Blomstrom for assistance with animal training and data collection and A. Cummins for preparing the histological material.

## Bibliography

Amengual, J. L., Di Bello, F., Ben Hadj Hassen, S., & Ben Hamed, S. (2022). Distractibility and impulsivity neural states are distinct from selective attention and modulate the implementation of spatial attention. Nature Communications, 13(1), 4796. 10.1038/s41467-022-32385-y

Awh, E., & Jonides, J. (2001). Overlapping mechanisms of attention and spatial working memory. Trends in Cognitive Sciences, 5(3), 119–126. 10.1016/S1364-6613(00)01593-X

Beauchamp, M. S., Petit, L., Ellmore, T. M., Ingeholm, J., & Haxby, J. V. (2001). A Parametric fMRI Study of Overt and Covert Shifts of Visuospatial Attention. NeuroImage, 14(2), 310–321. 10.1006/nimg.2001.0788

Beckers, G., Canavan, A. G. M., Zangemeister, W. H., & Homberg, V. (1992). Transcranial magnetic stimulation of human frontal and parietal cortex impairs programming of periodic saccades. NeuroOphthalmology, 12(5), 289–295. 10.3109/01658109209036984

Bowling, J. T., Friston, K. J., & Hopfinger, J. B. (2020). Top-down versus bottom-up attention differentially modulate frontal-parietal connectivity. Human Brain Mapping, 41(4), 928–942. 10.1002/hbm.24850

Chica, A. B., Bartolomeo, P., & Lupiáñez, J. (2013). Two cognitive and neural systems for endogenous and exogenous spatial attention. Behavioural Brain Research, 237, 107–123. 10.1016/j.bbr.2012.09.027

Corbetta, M. (1998). Frontoparietal cortical networks for directing attention and the eye to visual locations: Identical, independent, or overlapping neural systems? Proceedings of the National Academy of Sciences, 95(3), 831–838. 10.1073/pnas.95.3.831

De Haan, B., Morgan, P. S., & Rorden, C. (2008). Covert orienting of attention and overt eye movements activate identical brain regions. Brain Research, 1204, 102–111. 10.1016/j.brainres.2008.01.105

De Sousa, C., Gaillard, C., Di Bello, F., Ben Hadj Hassen, S., & Ben Hamed, S. (2021). Behavioral validation of novel high resolution attention decoding method from multi-units & local field potentials. NeuroImage, 231, 117853. 10.1016/j.neuroimage.2021.117853

Desimone, R., & Duncan, J. (1995). Neural Mechanisms of Selective Visual Attention. Annual Review of Neuroscience, 18(1), 193–222. 10.1146/annurev.ne.18.030195.001205

Di Bello, F., Ben Hadj Hassen, S., Astrand, E., & Ben Hamed, S. (2022). Prefrontal Control of Proactive and Reactive Mechanisms of Visual Suppression. Cerebral Cortex, 32(13), 2745–2761. 10.1093/cercor/bhab378

Dubey, A., Markowitz, D. A., & Pesaran, B. (2023). Top-down control of exogenous attentional selection is mediated by beta coherence in prefrontal cortex. Neuron, 111(20), 3321-3334.e5. 10.1016/j.neuron.2023.06.025

Fernández, A., Hanning, N. M., & Carrasco, M. (2023). Transcranial magnetic stimulation to frontal but not occipital cortex disrupts endogenous attention. Proceedings of the National Academy of Sciences of the United States of America, 120(10), e2219635120. 10.1073/pnas.2219635120

Fiebelkorn, I. C., & Kastner, S. (2020). Functional Specialization in the Attention Network. Annual Review of Psychology, 71(1), 221–249. 10.1146/annurev-psych-010418-103429

Folk, C. L., Remington, R. W., & Johnston, J. C. (1992). Involuntary covert orienting is contingent on attentional control settings. Journal of Experimental Psychology: Human Perception and Performance, 18(4), 1030–1044. 10.1037/0096-1523.18.4.1030

Friedman, N. P., & Robbins, T. W. (2022). The role of prefrontal cortex in cognitive control and executive function. Neuropsychopharmacology, 47(1), 72–89. 10.1038/s41386-021-01132-0

Gabay, S., Henik, A., & Gradstein, L. (2010). Ocular motor ability and covert attention in patients with Duane Retraction Syndrome. Neuropsychologia, 48(10), 3102–3109. 10.1016/j.neuropsychologia.2010.06.022

Gaillard, C., Ben Hadj Hassen, S., Di Bello, F., Bihan-Poudec, Y., VanRullen, R., & Ben Hamed, S. (2020). Prefrontal attentional saccades explore space rhythmically. Nature Communications, 11(1), 925. 10.1038/s41467-020-14649-7

Genovesio, A., Seitz, L. K., Tsujimoto, S., & Wise, S. P. (2016). Context-Dependent Duration Signals in the Primate Prefrontal Cortex. Cerebral Cortex, 26(8), 3345–3356. 10.1093/cercor/bhv156

Grosbras, M.-H., & Paus, T. (2002). Transcranial Magnetic Stimulation of the Human Frontal Eye Field: Effects on Visual Perception and Attention. Journal of Cognitive Neuroscience, 14(7), 1109–1120. 10.1162/089892902320474553

Hunt, A. R., Reuther, J., Hilchey, M. D., & Klein, R. M. (2019). The Relationship Between Spatial Attention and Eye Movements. In T. Hodgson (Ed.), Processes of Visuospatial Attention and Working Memory (Vol. 41, pp. 255–278). Springer International Publishing. 10.1007/7854_2019_95

Juan, C.-H., Shorter-Jacobi, S. M., & Schall, J. D. (2004). Dissociation of spatial attention and saccade preparation. Proceedings of the National Academy of Sciences of the United States of America, 101(43), 15541–15544. 10.1073/pnas.0403507101

Keefe, J. M., Pokta, E., & Störmer, V. S. (2021). Cross-modal orienting of exogenous attention results in visual-cortical facilitation, not suppression. Scientific Reports, 11(1), 10237. 10.1038/s41598-021-89654-x

Kim, C., Johnson, N. F., Cilles, S. E., & Gold, B. T. (2011). Common and Distinct Mechanisms of Cognitive Flexibility in Prefrontal Cortex. The Journal of Neuroscience, 31(13), 4771–4779. 10.1523/JNEUROSCI.5923-10.2011

Klein R.M. (1980). Does oculomotor readiness mediate cognitive control of visual attention? In R. S. Nickerson (Ed.), Attention and Performance (0 ed., pp. 259–276). Psychology Press. 10.4324/9781315802961

Kowler, E., Anderson, E., Dosher, B., & Blaser, E. (1995). The role of attention in the programming of saccades. Vision Research, 35(13), 1897–1916. 10.1016/0042-6989(94)00279-U

Landry, M., Da Silva Castanheira, J., Sackur, J., & Raz, A. (2021). Investigating how the modularity of visuospatial attention shapes conscious perception using type I and type II signal detection theory. Journal of Experimental Psychology: Human Perception and Performance, 47(3), 402–422. 10.1037/xhp0000810

Lowet, E., Gomes, B., Srinivasan, K., Zhou, H., Schafer, R. J., & Desimone, R. (2018a). Enhanced Neural Processing by Covert Attention only during Microsaccades Directed toward the Attended Stimulus. Neuron, 99(1), 207-214.e3. 10.1016/j.neuron.2018.05.041

Lowet, E., Gomes, B., Srinivasan, K., Zhou, H., Schafer, R. J., & Desimone, R. (2018b). Enhanced Neural Processing by Covert Attention only during Microsaccades Directed toward the Attended Stimulus. Neuron, 99(1), 207-214.e3. 10.1016/j.neuron.2018.05.041

Luck, S. J., Gaspelin, N., Folk, C. L., Remington, R. W., & Theeuwes, J. (2021). Progress toward resolving the attentional capture debate. Visual Cognition, 29(1), 1–21. 10.1080/13506285.2020.1848949

Marcos, E., & Genovesio, A. (2016). Determining Monkey Free Choice Long before the Choice Is Made: The Principal Role of Prefrontal Neurons Involved in Both Decision and Motor Processes. Frontiers in Neural Circuits, 10. 10.3389/fncir.2016.00075

Martinez-Trujillo, J. (2022). Visual Attention in the Prefrontal Cortex. Annual Review of Vision Science, 8(Volume 8, 2022), 407–425. 10.1146/annurev-vision-100720-031711

Messinger, A., Cirillo, R., Wise, S. P., & Genovesio, A. (2021). Separable neuronal contributions to covertly attended locations and movement goals in macaque frontal cortex. Science Advances, 7(14), eabe0716. 10.1126/sciadv.abe0716

Meyer, K. N., Du, F., Parks, E., & Hopfinger, J. B. (2018). Exogenous vs. endogenous attention: Shifting the balance of fronto-parietal activity. Neuropsychologia, 111, 307–316. 10.1016/j.neuropsychologia.2018.02.006

Michalczyk, Ł., Bielas, J., & Schab, A. (2020). Preparation of saccade sequences and eye programming affect endogenous covert attention. European Journal of Neuroscience, 52(5), 3419–3433. 10.1111/ejn.14773

Moore, T., & Armstrong, K. M. (2003). Selective gating of visual signals by microstimulation of frontal cortex. Nature, 421(6921), 370–373. 10.1038/nature01341

Moore, T., Armstrong, K. M., & Fallah, M. (2003). Visuomotor Origins of Covert Spatial Attention. Neuron, 40(4), 671–683. 10.1016/S0896-6273(03)00716-5

Moore, T., & Fallah, M. (2001). Control of eye movements and spatial attention. Proceedings of the National Academy of Sciences, 98(3), 1273–1276. 10.1073/pnas.98.3.1273

Müller, H. J., & Rabbitt, P. M. (1989). Reflexive and voluntary orienting of visual attention: Time course of activation and resistance to interruption. Journal of Experimental Psychology. Human Perception and Performance, 15(2), 315–330. 10.1037//0096-1523.15.2.315

Muri, R. M., Vermersch, A. I., Rivaud, S., Gaymard, B., & Pierrot-Deseilligny, C. (1996). Effects of single-pulse transcranial magnetic stimulation over the prefrontal and posterior parietal cortices during memory-guided saccades in humans. Journal of Neurophysiology, 76(3), 2102–2106. 10.1152/jn.1996.76.3.2102

Murthy, A., Thompson, K. G., & Schall, J. D. (2001). Dynamic Dissociation of Visual Selection From Saccade Programming in Frontal Eye Field. Journal of Neurophysiology, 86(5), 2634–2637. 10.1152/jn.2001.86.5.2634

Nobre, A. C., Gitelman, D. R., Dias, E. C., & Mesulam, M. M. (2000). Covert Visual Spatial Orienting and Saccades: Overlapping Neural Systems. NeuroImage, 11(3), 210–216. 10.1006/nimg.2000.0539

Panichello, M. F., & Buschman, T. J. (2021). Shared mechanisms underlie the control of working memory and attention. Nature, 592(7855), 601–605. 10.1038/s41586-021-03390-w

Perry, R. J., & Zeki, S. (2000). The neurology of saccades and covert shifts in spatial attention. Brain, 123(11), 2273–2288. 10.1093/brain/123.11.2273

Pinto, Y., Van Der Leij, A. R., Sligte, I. G., Lamme, V. A. F., & Scholte, H. S. (2013). Bottom-up and top-down attention are independent. Journal of Vision, 13(3), 16–16. 10.1167/13.3.16

Rafal, R. D., Posner, M. I., Friedman, J. H., Inhoff, A. W., & Bernstein, E. (1988). ORIENTING OF VISUAL ATTENTION IN PROGRESSIVE SUPRANUCLEAR PALSY. Brain, 111(2), 267–280. 10.1093/brain/111.2.267

Reynolds, J. H., & Heeger, D. J. (2009). The Normalization Model of Attention. Neuron, 61(2), 168–185. 10.1016/j.neuron.2009.01.002

Rigotti, M., Barak, O., Warden, M. R., Wang, X.-J., Daw, N. D., Miller, E. K., & Fusi, S. (2013). The importance of mixed selectivity in complex cognitive tasks. Nature, 497(7451), 585–590. 10.1038/nature12160

Rizzolatti, G., Riggio, L., Dascola, I., & Umiltá, C. (1987). Reorienting attention across the horizontal and vertical meridians: Evidence in favor of a premotor theory of attention. Neuropsychologia, 25(1), 31–40. 10.1016/0028-3932(87)90041-8

Rougier, N. P., Noelle, D. C., Braver, T. S., Cohen, J. D., & O’Reilly, R. C. (2005). Prefrontal cortex and flexible cognitive control: Rules without symbols. Proceedings of the National Academy of Sciences, 102(20), 7338–7343. 10.1073/pnas.0502455102

Sapountzis, P., Paneri, S., Papadopoulos, S., & Gregoriou, G. G. (2022). Dynamic and stable population coding of attentional instructions coexist in the prefrontal cortex. Proceedings of the National Academy of Sciences, 119(40), e2202564119. 10.1073/pnas.2202564119

Sato, T. R., & Schall, J. D. (2003). Effects of Stimulus-Response Compatibility on Neural Selection in Frontal Eye Field. Neuron, 38(4), 637–648. 10.1016/S0896-6273(03)00237-X

Sereno, A. B., Briand, K. A., Amador, S. C., & Szapiel, S. V. (2006). Disruption of Reflexive Attention and Eye Movements in an Individual with a Collicular Lesion. Journal of Clinical and Experimental Neuropsychology, 28(1), 145–166. 10.1080/13803390590929298

Sheliga, B. M., Riggio, L., & Rizzolatti, G. (1994). Orienting of attention and eye movements. Experimental Brain Research, 98(3). 10.1007/BF00233988

Smith, D. T., Rorden, C., & Jackson, S. R. (2004). Exogenous Orienting of Attention Depends upon the Ability to Execute Eye Movements. Current Biology, 14(9), 792–795. 10.1016/j.cub.2004.04.035

Smyth, M. M., & Scholey, K. A. (1994). Interference in immediate spatial memory. Memory & Cognition, 22(1), 1–13. 10.3758/BF03202756

Tehovnik, E. J. (1996). Electrical stimulation of neural tissue to evoke behavioral responses. Journal of Neuroscience Methods, 65(1), 1–17. 10.1016/0165-0270(95)00131-x

Theeuwes, J. (1992). Perceptual selectivity for color and form. Perception & Psychophysics, 51(6), 599–606. 10.3758/BF03211656

Theeuwes, J. (2014). Spatial orienting and attentional capture. In A. C. (Kia) Nobre & S. Kastner (Eds.), The Oxford Handbook of Attention (Vol. 1). Oxford University Press. 10.1093/oxfordhb/9780199675111.001.0001

Thickbroom, G. W., Stell, R., & Mastaglia, F. L. (1996). Transcranial magnetic stimulation of the human frontal eye field. Journal of the Neurological Sciences, 144(1–2), 114–118. 10.1016/S0022-510X(96)00194-3

Thompson, K. G., Bichot, N. P., & Schall, J. D. (1997). Dissociation of Visual Discrimination From Saccade Programming in Macaque Frontal Eye Field. Journal of Neurophysiology, 77(2), 1046–1050. 10.1152/jn.1997.77.2.1046

Thompson, K. G., Biscoe, K. L., & Sato, T. R. (2005). Neuronal Basis of Covert Spatial Attention in the Frontal Eye Field. The Journal of Neuroscience, 25(41), 9479–9487. 10.1523/JNEUROSCI.0741-05.2005

Xia, R., Chen, X., Engel, T. A., & Moore, T. (2024). Common and distinct neural mechanisms of attention. Trends in Cognitive Sciences, 28(6), 554–567. 10.1016/j.tics.2024.01.005

