## SUPPLEMENTARY MATERIALS for "Dynamic Attentional Control Through Mixed Prefrontal Cortex Resources"

| pre GO | post GO | post Saccade |
|---|---|---|
| Motor 15% (n = 35) | Motor 34.3% (n = 12)<br>Attention 8.6% (n = 3)<br>Both 5.7% (n = 2)<br>Untuned 51.4% (n = 18) | Motor 51.4% (n = 18)<br>Attention 0%<br>Both 5.7% (n = 2)<br>Untuned 42.8% (n = 15) |
| Attention 9% (n = 22) | Motor 4.5% (n = 1)<br>Attention 50.0% (n = 11)<br>Both 0%<br>Untuned 45.5% (n = 10) | Motor 40.9% (n = 9)<br>Attention 9.1% (n = 2)<br>Both 9.1% (n = 2)<br>Untuned 40.9% (n = 9) |
| Both 4% (n = 9) | Motor 22.2% (n = 2)<br>Attention 44.4% (n = 4)<br>Both 11.1% (n = 1)<br>Untuned 22.2% (n = 2) | Motor 66.7% (n = 6)<br>Attention 0%<br>Both 0%<br>Untuned 33.3% (n = 3) |
| Untuned 72% (n = 172) | Motor 7.6% (n = 13)<br>Attention 9.3% (n = 16)<br>Both 4.6% (n = 8)<br>Untuned 78.5% (n = 135) | Motor 27.9% (n = 48)<br>Attention 6.4% (n = 11)<br>Both 8.1% (n = 14)<br>Untuned 57.6% (n = 99) |

**Table S1. Unit selectivity evolved across task epochs.** Number and relative proportion of Motor, Attention, Both, and untuned units in the Pre-Go epoch of cued trials (1st column). Number and relative proportion of the same units in the period following the Go signal (post Go, 2$^{nd}$ column) and the saccadic response (post Saccade, 3$^{rd}$ column), grouped by the unit's initial (pre Go) selectivity. See Fig. 2C legend for epoch definitions.

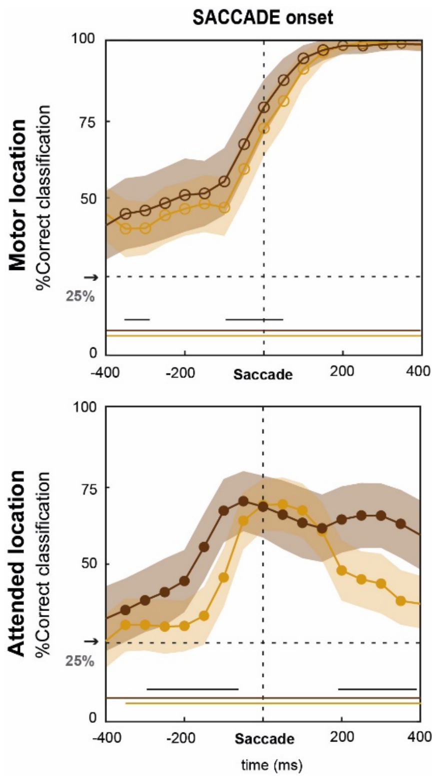


**Figure S1**. **Population decoding relative to saccade onset.** Time course of accuracy for decoding the Motor Location (top) and Attended Location (bottom) for cued (dark brown) and uncued (light brown) trials, aligned to saccade onset. Lines and shading show the mean classification accuracy ± s.e. in each sliding window. Black horizontal lines indicate a significant difference between accuracy on cued and uncued trials ($p < 0.001$). Colored lines indicate periods of significant ($p < 0.001$) above-chance classification for cued (dark brown) and uncued (light brown) trials.

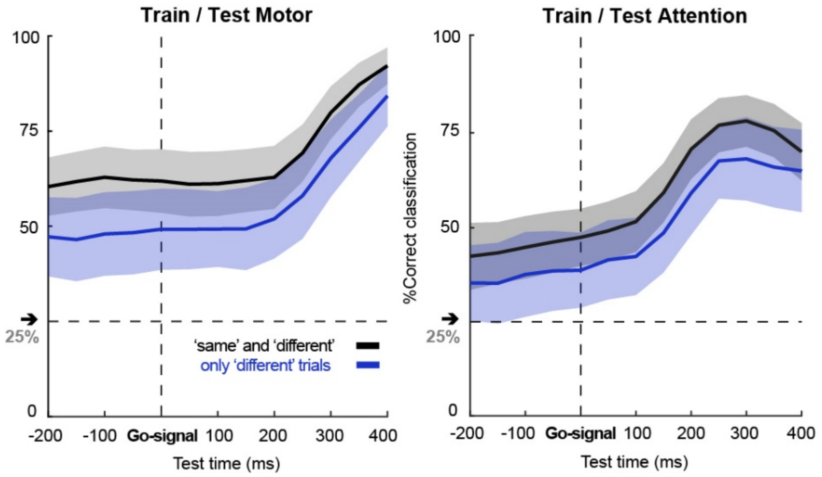


**Figure S2. Decoding accuracy for 'same' and 'different' location trials.** Average within-modality classification accuracies were calculated using just 'different' location trials (blue) and using both 'same' and 'different' location trials (black). Curves show accuracies along the diagonal for the cross-temporal classification analyses, aligned to the Go-signal. Classification of both the Motor and Attended Locations was better when 'same' location trials were included and only this single location had to be covertly monitored and targeted.

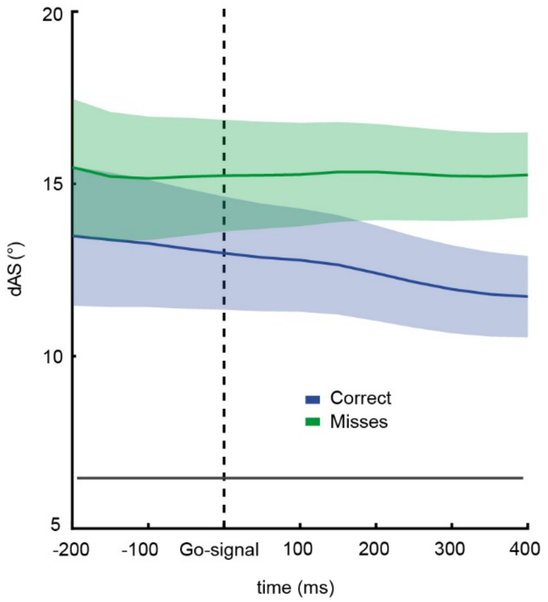


**Figure S3. Attention localization as a function of behavioral success.** Displacement of the attentional spotlight (dAS) calculated for correct and saccade omission (Miss) trials, aligned to the Go-signal. The black line at the bottom indicates that, for all time points shown, the locus of attention was significantly ($p < 0.001$) further from the Go-signal location (attention target) when the Go-signal went undetected (Miss trials) than when it was detected (Correct trials).

### Supplemental information

#### *Supplementary Note 1: motor and attention accuracies dynamic using both 'same' and 'different' trials*

Both the decoding accuracies consistently exceeded chance level (25%) in the epoch preceding the Go-signal presentation ($p < 0.001$, for all time bins between -200 and 0, Go-signal aligned), and reached approximately 60% for testing motor variables and 47% for testing attention variables at Go-signal onset. Following the presentation of the Go-signal, both the decoding performances steadily increased, with motor predictions reaching their higher value at 400 ms (i.e., at the end of the time window considered) while attention showed its peak at around 350 ms (~79%).
